# Comparing statistical learning methods for complex trait prediction from gene expression

**DOI:** 10.1101/2024.06.01.596951

**Authors:** Noah Klimkowski Arango, Fabio Morgante

## Abstract

Accurate prediction of complex traits is an important task in quantitative genetics that has become increasingly relevant for personalized medicine. Genotypes have traditionally been used for trait prediction using a variety of methods such as mixed models, Bayesian methods, penalized regressions, dimension reductions, and machine learning methods. Recent studies have shown that gene expression levels can produce higher prediction accuracy than genotypes. However, only a few prediction methods were used in these studies. Thus, a comprehensive assessment of methods is needed to fully evaluate the potential of gene expression as a predictor of complex trait phenotypes. Here, we used data from the *Drosophila* Genetic Reference Panel (DGRP) to compare the ability of several existing statistical learning methods to predict starvation resistance from gene expression in the two sexes separately. The methods considered differ in assumptions about the distribution of gene effect sizes – ranging from models that assume that every gene affects the trait to more sparse models – and their ability to capture gene-gene interactions. We also used functional annotation (*i*.*e*., Gene Ontology (GO)) as an external source of biological information to inform prediction models. The results show that differences in prediction accuracy between methods exist, although they are generally not large. Methods performing variable selection gave higher accuracy in females while methods assuming a more polygenic architecture performed better in males. Incorporating GO annotations further improved prediction accuracy for a few GO terms of biological significance. Biological significance extended to the genes underlying highly predictive GO terms with different genes emerging between sexes. Notably, the Insulin-like Receptor (*InR*) was prevalent across methods and sexes. Our results confirmed the potential of transcriptomic prediction and highlighted the importance of selecting appropriate methods and strategies in order to achieve accurate predictions.

## Introduction

Predicting yet-to-be observed phenotypes for complex traits is an important task for many branches of quantitative genetics. Complex trait prediction was developed in agricultural breeding to select the best performing individuals for economically important traits such as milk yield in dairy cattle using estimated breeding values (EBVs). While EBVs have been traditionally computed using pedigree information, with the availability of genotyping arrays, EBVs have been replaced or supplemented with their genomic counterpart – genomic EBVs (GEBVs) [1, 2]. GEBVs are linear combinations of the genotypes of the target individuals and the effect sizes of many genetic variants along the genome computed in a reference population. The same concept has later been applied to human genetics, especially in the context of precision medicine. Here the goal is to predict medically relevant phenotypes such as body mass index (BMI) or disease susceptibility using Polygenic Scores (PGSs) [3, 4]. While GEBVs and PGSs are technically the same, the different goals of these two quantities (*i*.*e*., selection for GEBVs and prevention/monitoring for PGS) have important implications. We refer the readers to [2] for a comprehensive treatment of this topic.

The estimation of the effect sizes of genetic variants to be used for prediction can be done using a variety of methods. The most common methods have regression at their core, where the response variable is the phenotype of interest and the predictor variables are the genotypes [5]. Because the number of genetic variants is usually much larger than the sample size, methods that perform some regularization of the effect sizes are needed. These methods encompass dimension reduction methods (*e*.*g*., principal components regression), penalized regression methods (*e*.*g*., ridge regression), linear mixed models (*e*.*g*., GBLUP), Bayesian methods (*e*.*g*., BayesC), and machine learning (*e*.*g*., random forest) [6–10]. These methods differ in the assumptions they make regarding the distribution of the effect sizes, with some methods performing only effect shrinkage and some methods performing variable selection as well [5]. Research focused on comparing several methods has shown that there is no single best method, with performance being affected by the genetic architecture of the trait of interest (*e*.*g*., sparse vs dense), the biology of the species (*e*.*g*., the extent of linkage disequilibrium), and assumptions of the method [11, 12].

Traditionally, genotype data has been used for complex trait prediction since they are easy and cost-effective to obtain. However, it is now possible to obtain multidimensional molecular data such as gene expression or metabolite levels at a reasonable cost. This has opened to the possibility of using these additional layers of data for complex trait prediction. Given that genetic information flows from DNA to RNA, to proteins, to metabolites to affect phenotypes [13], using these intermediate layers of data could improve prediction accuracy for at least some traits. In addition, to being biologically ‘closer’ to phenotypes, gene expression levels, protein levels, and metabolite levels can be thought of as endophenotypes, which are affected by environmental conditions as well. Thus, endophenotypes could capture environmental and gene-by-environment effects [14]. Recent work has shown that using additional omic data types can result in more accurate predictions [14–19]. In particular, using transcriptomic data has shown good promise. For example, Wheeler *et al*. used lymphoblastoid cell line data to show that gene expression levels provided much higher accuracy than genotypes at predicting intrinsic growth rate [15]. In a similar fashion, Morgante *et al*. found that prediction accuracy of starvation resistance in *Drosophila melanogaster* was higher when using gene expression levels than when using genotypes [17].

While these studies have shown the potential of gene expression as a predictor of complex phenotypes, only a few statistical methods were used, with most studies using linear mixed models. However, as discussed above, studies using genotypes have found that prediction accuracy can vary substantially depending on the method used. Thus, in this study we sought to compare several common methods spanning dimensionality reductions, penalized linear regressions, Bayesian linear regressions, linear mixed models, and machine learning in their ability to predict starvation resistance from gene expression levels using data from the *Drosophila* Genetic Reference Panel [20].

## Materials and Methods

### Data Processing

The Drosophila melanogaster Genetic Reference Panel (DGRP) is a collection of over 200 inbred lines derived from a natural population that have full genome sequences and phenotypic measurements for several traits [21]. Additionally, prior work from [22] obtained full transcriptome profiles by RNA sequencing for a total of 200 DGRP lines. Following the filtering steps described in [17], we ended up with 11,338 genetically variable and highly expressed genes in females and 13,575 in males. Among the many traits available for the DGRP, in this work we used starvation resistance as a model trait because it can be predicted with decent accuracy with the small sample size available [17]. In particular, line means for 198 lines that have both transcriptome profiles and phenotypic measurements, adjusted for the effect of *Wolbachia* infection and major inversions [20] were used for all the analyses.

### Transcriptomic Prediction

The methods used follow the general multiple regression model:

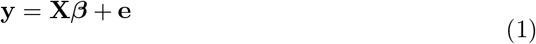

where **y** is a *n*-vector of phenotypes, **X** is a *n* × *m* matrix of expression levels for *m* genes, ***β*** is a *m*-vector of effect sizes, and **e** is a *n*-vector of residuals. We assume that the columns of **X** and **y** have been centered to mean 0.

### Principal Component Regression (PCR)

PCR [23] uses Principal Component Analysis [24] to reduce the dimensionality of the predictor matrix, **X**, by selecting a set of *k* orthogonal components that are linear combinations of the original predictors and maximize their variance. Then, the *n* × *k* matrix of principal components, **T**, is used in place of **X** in equation 1 [25]. We used the algorithm implemented in the R package pls v. 2.8-2 ( [26]) with default parameters. We used 5-fold cross validation in the training set to select the number of principal components to be used for prediction in the test set.

### Partial Least Squares Regression (PLSR)

Like PCR, PLSR [27] also reduces the dimensionality of the predictor matrix, **X**. However, this is achieved via a simultaneous decomposition of **X** and **y** and selecting a set of *k* components that maximizes the covariance between **X** and **y**. This addresses a limitation of PCR that the components that best “explain” **X** may not necessarily be the most relevant to **Y**. Then, the *n* × *k* matrix of latent vectors, **T**, is used in place of **X** in equation 1 [28]. We used the algorithm implemented in the R package pls v. 2.8-2 ( [26]), setting the method to ‘widekernelpls’ which is suitable for the wide (*m >> n*) matrix of gene expression, and other default parameters. We used 5-fold cross validation in the training set to select the number of latent vectors to be used for prediction in the test set.

### Ridge Regression (RR)

Ridge Regression [6] is a penalized linear regression method that uses an *ℓ*_2_ penalty to achieve shrinkage of the estimates of the effect sizes. The amount of shrinkage is determined by a tuning parameter, *λ*, such that large values of *λ* result in more shrinkage. We used the algorithm implemented in the R package glmnet v. 4.1-8 ( [29]). We used 5-fold cross validation to select *λ*. All other parameters were left as their default values.

### Least Absolute Shrinkage Selector Operator (LASSO)

LASSO [30] is a penalized linear regression method that uses an *ℓ*_1_ penalty to perform both variable selection (by setting some effects to be exactly 0) and shrinkage of the estimates of the effect sizes. The amount of shrinkage and variable selection is determined by a tuning parameter, *λ*, such that large values of *λ* result in more shrinkage. We used the algorithm implemented in the R package glmnet v. 4.1-8 ( [29]) was fit. We used 5-fold cross validation to select *λ*. All other parameters were left as their default values.

### BayesC

BayesC [31] is a Bayesian linear regression method that imposes a spike-and-slab prior on the effect sizes:

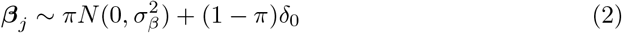

where *π* is the probability that the effect of *j*^*th*^ variable comes from a normal distribution with mean 0 and variance 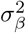 , and δ_0_ is a point-mass at 0. In this way, both variable selection and effect shrinkage are achieved. In the R package BGLR v. 1.1.0 implementation, [32], the posterior distribution of the effect sizes and some model parameters are computed using Markov Chain Monte Carlo (MCMC) methods. We ran the algorithm for 130,000 iterations, with the first 30,000 iterations discarded as burn-in and retaining every 50^*th*^ sample. We assessed convergence through visual inspection of the trace plots. The expected proportion of variance by the predictors, *R*^2^, was set to 0.8 to in line with the broad sense heritability of starvation resistance [33].

### Variational Bayesian Variable Selection (VARBVS)

VARBVS is a Bayesian linear regression method that imposes the same spike-and-slab prior as BayesC. However, posterior computations are done using Variational Inference rather than MCMC, which is more computationally efficient [34]. The algorithm implemented in the R package varbvs v. 2.6-10 ( [34]) was fit with default parameters.

### Multiple Regression with Adaptive Shrinkage (MR.ASH)

MR.ASH is a Bayesian linear regression method that imposes a scale mixture-of-normals prior on the effect sizes:

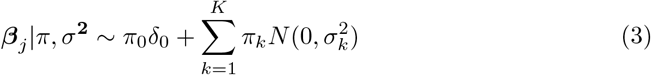

for a fixed grid of variances, *σ*^**2**^. Thus, like BayesC and VARBVS, MR.ASH performs both variable selection and effect shrinkage, but is able to model more complex distributions of the effect sizes owing to a more flexible prior. MR.ASH uses a Variational Empirical Bayes approach to estimate the prior (*i*.*e*., the mixture weights, *π*) from the data and compute the posterior distribution of the effect sizes [35]. The algorithm implemented in the R package mr.ash.alpha v. 0.1-43 ( [36]) was fit with default parameters, but used the effect size estimates from LASSO to initialize MR.ASH.

### Transcriptomic Best Linear Unbiased Predictor (TBLUP)

TBLUP [37] is a linear mixed model that aggregates the effects of all the genes into a single random effect. Let 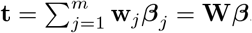, where **w**_*j*_ is a standardized version of **x**_*j*_ to have variance 1, then:

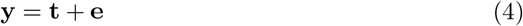

where **t** is a *n*-vector of transcriptomic effects, 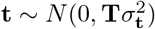, and 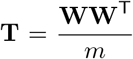 is the Transcriptomic Relationship Matrix (TRM). TBLUP was implemented by using theR package BGLR v. 1.1.0 ( [32]). BGLR uses a Bayesian approach to estimate the transcriptomic and residual variance components. We ran the algorithm for 85,000 iterations, with the first 10,000 iterations discarded as burn-in and retaining every 50^*th*^ sample. We assessed convergence through visual inspection of the trace plots. The expected proportion of variance, *R*^2^, was set to 0.8. All other parameters were left as their default values.

All the previous methods assume that no gene-gene interactions affect starvation resistance. Thus, we decided to add some more flexible machine learning methods to the comparison, which are also able to capture interaction effects.

### Random Forest (RF)

Random Forest is a machine learning method whereby a collection of decision trees are grown, each on a different bootstrap sample of the predictor data [10]. This method has been used to identify gene-gene interactions successfully [38]. The model is given by:

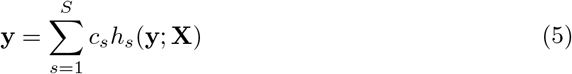

where *S* is the number of decision trees, *c*_*s*_ is a shrinkage factor that averages the trees, *h*_*s*_(**y**; **X**) is a decision tree that is grown using only a subset of predictors at each nodes [10]. The algorithm implemented in the R package partykit v. 1.2-20 ( [39]) was fit with 1000 trees and default parameters.

### Neural Network (NN)

Artificial Neural Networks are a type of machine learning method that use layers of nodes, or neurons, to process data similar to how the human brain works. Neural networks are built using input layers, hidden layers, and output layers. Input layers receive data while output layers calculate final results. Hidden layers are able to modify inputs from previous layers to discover trends or patterns within data [40]. Network assembly is tailored to individual problems, as networks can vary by hidden layer count, neuron count per layer, and activation function per layer. In our model, weights for a single layer of hidden neurons were estimated using resilient backpropagation [41].

Neural networks use nonlinear activation functions to determine whether neurons in hidden layers should be activated based on their inputs. This feature can be used to model gene interactions [42]. The general model for an artificial neural network is given by

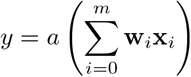

where *y* is the phenotype of a given line, *a* is an activation function, *m* is the number of inputs, **w** is the vector of weights, **x** is a vector of gene expression, and *w*_0_ is the bias or intercept of the model. The neural network implemented through the R package neuralnet v1.44.2 ( [43]) used default parameters and a custom neuron structure. Neuron count selection is a fundamental problem in constructing networks [44]. We specified a hidden layer size of 1,000 neurons.

### Gene Ontology Informed Transcriptomic Prediction

While some of the methods above try to enrich the prediction model for genes that are particularly predictive of the trait by performing internal variable selection, this procedure becomes difficult with small sample size such as in the DGRP. Informing prediction models with functional annotation has been shown to be effective at disentangling signal from noise and improve accuracy in complex trait prediction [17, 45–47]. Edwards *et al*. (2016) and Morgante *et al*. (2020) used Gene Ontology (GO) [48] to improve prediction accuracy for three complex traits in *Drosophila* [17, 45]. However, these applications only used BLUP-type models to include GO information. Here, we tested a few additional methods described below. For each sex, we selected GO terms that included at least five genes present in the DGRP expression data as done in previous work [17]. This procedure resulted in 2,628 terms for females and 2,580 terms for males being retained for further analysis. For all methods, GO-informed models were fit with one GO term at a time for all GO terms specified for each sex.

### Sparse Group LASSO

Sparse group LASSO [49] is a penalized regression method that uses a combination of the *ℓ*_1_ penalty and a group LASSO penalty [50]. The Group LASSO [51] applies variable selection on entire groups of predictors, while the *ℓ*_1_ penalty achieves effect shrinkage and variable selection at the individual variable level. The strength of the penalties is determined by a tuning parameter, *λ*, such that larger values of *λ* result in more shrinkage/selection. In our application, one group included all the genes in the selected GO term and the other group included all the remaining genes. We used the Sparse Group LASSO implementation in the `R` package `sparseGL` v1.0.2 [52] with default parameters.

### GO-BayesC

GO-BayesC is a Bayesian linear regression method that imposes independent spike-and-slab priors on effect sizes of genes grouped by GO term association. Let

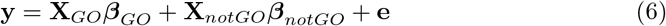

where **X**_*GO*_ is the subset of **X** containing the genes associated with the selected GO term, ***β***_GO_ is the vector of effects of the genes in the selected GO term, **X**_*notGO*_ is the subset of **X** containing all other genes, and *β*_*notGO*_ the vector of effects of all other genes. *β*_*GO*,*j*_ and ***β***_notGO,j_ are assigned separate spike-and-slab prior distributions as in equation 2. This method uses the same algorithm from the `R` package `BGLR` [32]. We ran the algorithm for 130,000 iterations, with the first 30,000 iterations discarded as burn-in and retaining every 50^*th*^ sample. We assessed convergence through visual inspection of the trace plots. The expected proportion of variance by all the predictors, *R*^2^, was set to 0.8.

### GO-TBLUP

GO-TBLUP is an extension of TBLUP that includes two random effects, one associated with genes in the selected GO term and one associated with all the other genes:

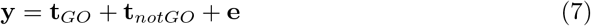

where **t**_*GO*_ is a *n*-vector of transcriptomic effects associated with genes in the GO term,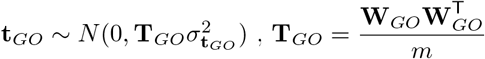, **W**_*GO*_ is the subset of **W** containing the genes associated with the selected GO^m^ term, **t**_*notGO*_ is a *n*-vector of transcriptomic effects associated with all other genes, 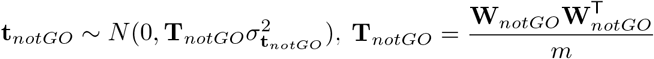, and **W**_*notGO*_ is the subset of **W** containing all other genes. GO-TBLUP was implemented by using the R package BGLR v. 1.1.0 ( [32]). BGLR uses a Bayesian approach to estimate the transcriptomic and residual variance components. We ran the algorithm for 85,000 iterations, with the first 10,000 iterations discarded as burn-in and retaining every 50^*th*^ sample. We assessed convergence through visual inspection of the trace plots. The expected proportion of variance, *R*^2^, was set to 0.8. All other parameters were left as their default values.

### Evaluation Scheme

We fitted each method to 90% of the data (*i*.*e*., training set) to estimate the model parameters and used them to predict phenotypes for the remaining 10% of the data (*i*.*e*., test set). Prediction accuracy was measured as the correlation coefficient between actual and predicted phenotypes. We repeated this procedure for 25 random training-test splits and used the average correlation across splits as our final metric to assess prediction accuracy.

## Results

### Transcriptomic Prediction

We first fitted a few widely used regression methods and compared their prediction accuracy. The results are shown in Fig 1 and S1 Table.. Overall, prediction accuracy was moderately low, especially considering that the analyses were based on lines means of many individual flies, resulting in the majority of the phenotypic variance being genetic [17]. However, the results show that differences in prediction accuracy between methods exist, both within sex and across sexes. In males, we found that TBLUP (mean *r* = 0.455 ± 0.035) and Ridge Regression (*r* = 0.455 ± 0.034) provided the highest accuracy, with BayesC (mean *r* = 0.448 ± 0.033), PLSR (mean *r* = 0.442 ± 0.034), and PCR (mean *r* = 0.430 ± 0.039) being competitive. On the other hand, Neural Network provided the lowest prediction accuracy for males (*r* = 0.143 ± 0.079). In females, we observed more marked differences in prediction accuracy across methods. Methods that perform variable selection – *i*.*e*., VARBVS, MR.ASH, and LASSO – tended to perform better than the other methods, with VARBVS (mean *r* = 0.503 ± 0.031) providing the highest accuracy. Neural Network provided the lowest prediction accuracy in females (*r* = 0.064 *±* 0.051) as well.

**Fig 1.**
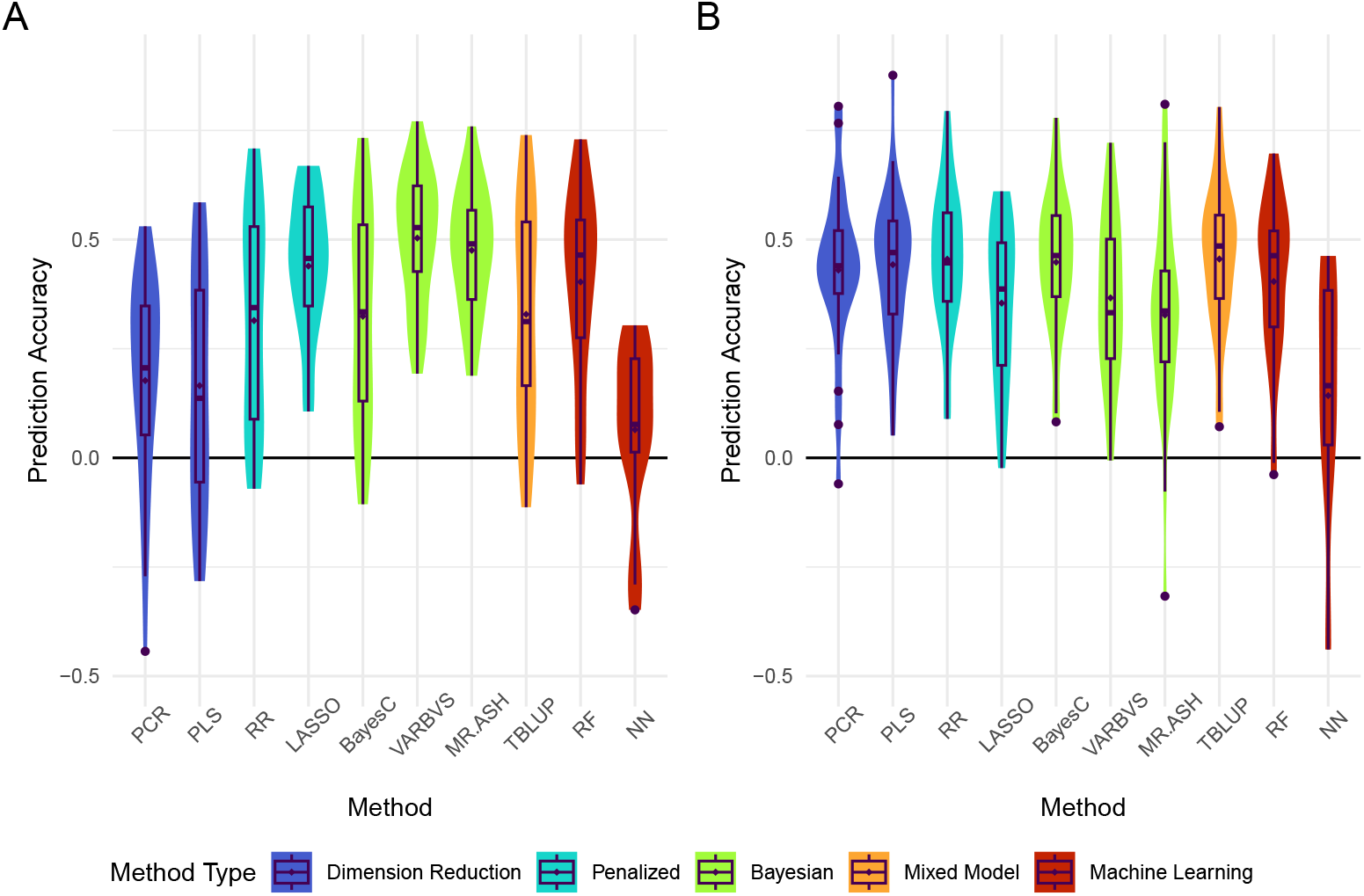
Prediction accuracy of 25 replicates in A) females and B) males for all standard methods. Methods are colored by family. The mean correlation coefficient is denoted by diamonds. Outliers are denoted by circles.

### Gene Ontology Informed Transcriptomic Prediction

It has been shown previously that informing prediction models with functional information can help achieve higher prediction accuracy [17, 45–47]. Thus, we also tested methods that could include external information. In this work, we focused on GO annotation and extension of BayesC and TBLUP, namely GO-BayesC and GO-TBLUP. The results are summarized in Fig 2 and S2 Table.. We also sought to use the Sparse Group LASSO. However, in our initial testing, the prediction accuracies provided by that method were nearly identical for all GO terms tested (S1 Fig.). This pattern was also seen for GO terms found to be highly predictive in GO-BayesC and GO-TBLUP. Thus, we decided not to assess Sparse Group Lasso further.

**Fig 2.**
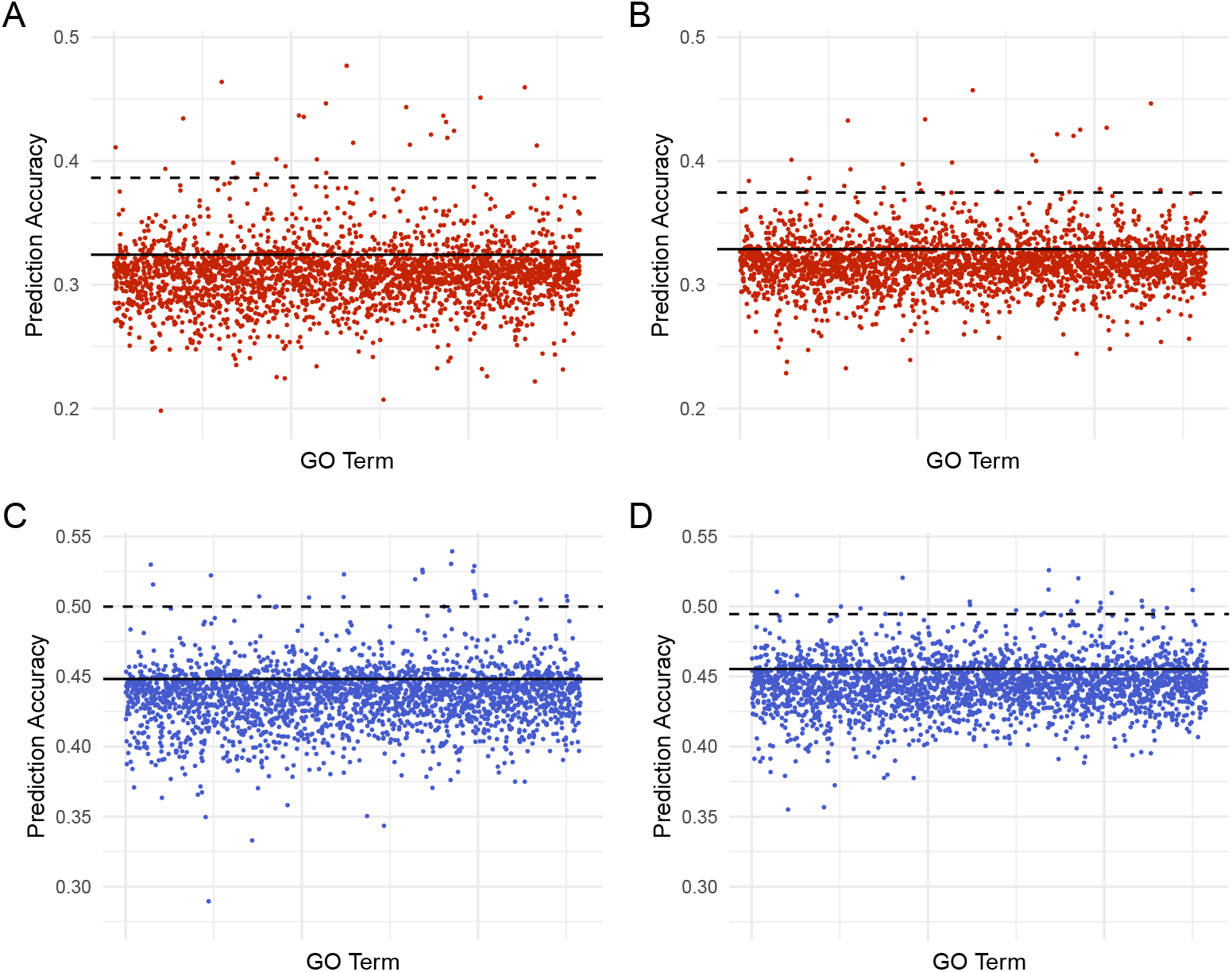
Prediction accuracy using GO-BayesC in females (A) and males (B). Prediction accuracy using GO-TBLUP in females (C) and males (D). Each dot represents the mean correlation between true and predicted phenotypes (*r*) across 25 replicates for a GO term. The solid line indicates the mean *r* from the respective standard method (*i*.*e*., BayesC and TBLUP). The dashed black line represents the 99^*th*^ percentile of terms ranked by prediction accuracy.

We found that GO-BayesC and GO-TBLUP provided accuracies that were similar to or lower than the respective standard model (*i*.*e*., BayesC and TBLUP) for the majority of GO terms in both sexes. However, some GO terms seemed to be particularly predictive of the trait, yielding accuracies that are substantially higher than the standard models. The accuracies provided by GO-BayesC and GO-TBLUP generally agreed (*r* = 0.836), as shown in Fig 3.

**Fig 3.**
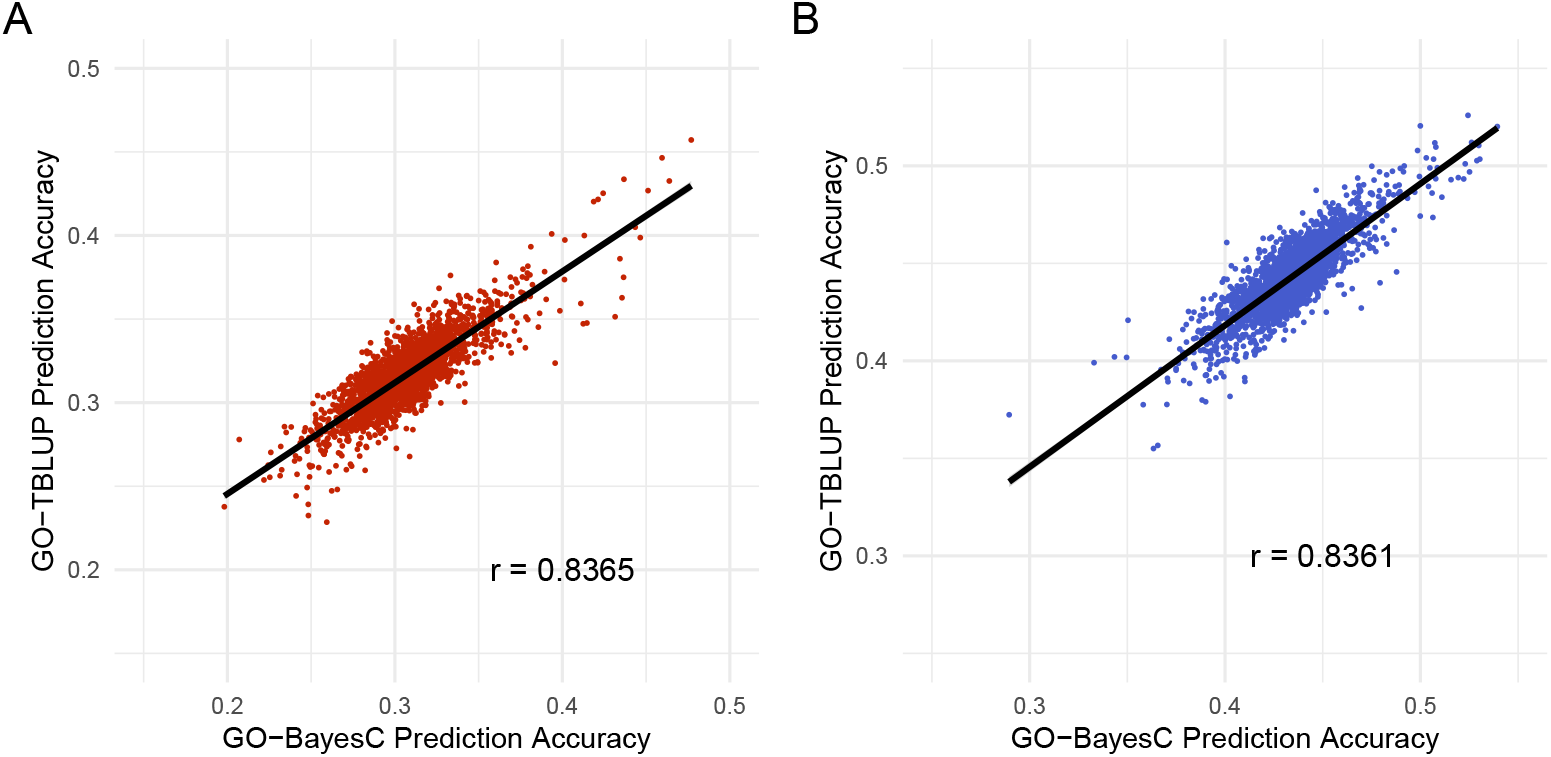
Plot of prediction accuracy for all GO terms using GO-BayesC(x-axis) against GO-TBLUP (y-axis) for A) females and B) males. The black line represents the line of best fit for each panel.

In females, four of the five most predictive GO terms are shared between GO-BayesC and GO-TBLUP. GO terms GO:0017056 (GO-BayesC *r* = 0.477 ± 0.034, GO-TBLUP *r* = 0.457 ± 0.041) and GO:0006606 (GO-BayesC *r* = 0464. ± 0.039, GO-TBLUP *r* = 0.434 ± 0.044) are both related to nuclear import by function and structure, respectively. Lee *et al*. has demonstrated starvation resistance inducing nuclear pore degradation in yeast [53]. The other two GO terms GO:0055088 (GO-BayesC *r* = 0.446 ± 0.040, GO-TBLUP *r* = 0.460 ± 0.036) and GO:0045819 (GO-BayesC *r* = 0.451 ± 0.038, GO-TBLUP *r* = 0.427 ± 0.042) are related to macromolecule metabolism in lipids and carbohydrates, respectively. GO:0055088 has been implicated in starvation resistance using Korean rockfish [54]. GO:0017056 was the most predictive term for GO-BayesC (*r* = 0.477 ± 0.034) and GO-TBLUP (*r* = 0.457 ± 0.041). However, some differences exist. For example, GO:0016042, which is involved with lipid catabolism, was found to be highly predictive by GO-BayesC (*r* = 0.447 ± 0.038), while GO:0008586, which is involved with wing vein morphogenesis, was highly predictive in GO-TBLUP (*r* = 0.424 ± 0.039). In males, two of the five most predictive GO terms are shared between methods. GO:0042593 (GO-BayesC *r* = 0539. ± 0.032, GO-TBLUP *r* = 0.520 ± 0.035) is involved in glucose homeostasis, while GO:0035003 (GO-BayesC *r* = 0.526 ± 0.036, GO-TBLUP *r* = 0.512 ± 0.030) is involved in the subapical complex. which is a key component of the intestinal epithelial tissue. As part of epithelial tissue, the subapical complex is involved with nutrient acquisition in the intestines as part of the barrier between host cells and the gut microbiome. [55] Four of the top five GO terms found from either method are implicated in cellular growth and development. In GO-BayesC, GO:0042461 (*r* = 0.530 ± 0.036) is involved in photoreceptor cells, GO:0001738 (*r* = 0.530 ± 0.036) is involved in epithelial tissue and GO:0045186 (*r* = 0.529 ± 0.037) is involved in the assembly of the zonula adherens. GO:0007485, which is involved in genital disc formation, was highly predictive in GO-TBLUP(*r* = 0.520 ± 0.030). Multiple studies have found connections between cell size regulation and overall body size with starvation resistance [56–58]. The top GO term for TBLUP in males GO:0035008(*r* = 0.526 ± 0.026 ), is involved in the positive regulation of the melanization defense response. The biological connection of this process to starvation resistance is unclear, as this response increases oxidative stress in wounds to prevent infection [59].

### Gene Analysis

Given that many of the most predictive GO terms were biologically relevant to starvation resistance, we investigated whether any particular genes were included in such terms. To do so, we selected the 1% most predictive GO terms for each method and sex (26 and 25 GO terms for females and males, respectively). For each prediction method and sex combination, we counted how many times each gene was found across these GO terms. We then examined the distribution of the count (S2 Fig.) and decided to focus only on the most frequently occurring genes. This resulted in selecting genes appearing in 5 or more GO terms for GO-TBLUP in females and genes appearing in 4 or more GO terms for all other method and sex combinations. These results are summarized in Table 1 and S3 Table..

**Table 1.** Top overlapping genes across the 1% most predictive GO terms for each method and sex combination. Genes appearing in all four setups are highlighted in green. Genes appearing in both methods for females are highlighted in red. Genes appearing for both methods in males are highlighted in blue.

| GO-BayesC Female |  | GO-TBLUP Female |  | GO-BayesC Male |  | GO-TBLUP Male |  |
| --- | --- | --- | --- | --- | --- | --- | --- |
| Gene | Count | Gene | Count | Gene | Count | Gene | Count |
| *AkhR* | 9 | *Egfr* | 8 | *sdt* | 13 | *sdt* | 8 |
| *mbo* | 6 | *Akt1* | 6 | *aPKC* | 9 | *aPKC* | 7 |
| *InR* | 6 | *AkhR* | 6 | *par-6* | 8 | *PDZ-GEF* | 6 |
| *Nup54* | 5 | *InR* | 6 | *Patj* | 8 | *par-6* | 6 |
| *Akh* | 5 | *Sik3* | 5 | *scrib* | 5 | *Patj* | 5 |
| *Nup154* | 4 | *emc* | 5 | *baz* | 5 | *crb* | 5 |
| *Nup93-1* | 4 | *put* | 5 | *crb* | 5 | *Ilp2* | 4 |
| *Nup205* | 4 | *babo* | 5 | *Moe* | 5 | *Desat1* | 4 |
| *Nup93-2* | 4 | *Pdk1* | 5 | *PDZ-GEF* | 5 | *InR* | 4 |
| *Nup98-96* | 4 | *Ack* | 5 | *AkhR* | 4 | *Ras85D* | 4 |
| *Nup153* | 4 | *Pkc98E* | 5 | *Ilp2* | 4 |  |  |
| *Akt1* | 4 | *CG3216* | 5 | *InR* | 4 |  |  |
| *Pi3K92E* | 4 | *CG31183* | 5 | *dlg1* | 4 |  |  |
| *Egfr* | 4 | *Erk7* | 5 | *shg* | 4 |  |  |
| *Erk7* | 4 | *hpo* | 5 |  |  |  |  |
|  |  | *hep* | 5 |  |  |  |  |

For both sexes, the results show that some overlapping genes were found by both GO-BayesC and GO-TBLUP while some genes were picked up by only one method. One trend that emerged across all setups a significant enrichment (Fisher’s Exact Test *P <* 0.001) of protein kinases among the top genes.

In females, GO-BayesC and GO-TBLUP found 5 genes in common out of the top 1% of GO terms. These five genes are related to insulin signaling and lipid metabolism. Insulin signaling is involved in cell growth, feeding, carbohydrate metabolism, and many other traits critical for survival [60]. The insulin-like receptor (*InR*) is crucial for insulin signaling in carbohydrate metabolism. The adipokinetic hormone receptor (*AkhR*) is responsible for both carbohydrate and lipid metabolism signals. *AkhR* was shown to coexpress with *InR* on starvation-induced hyperactivity [61]. *Akt1* is the core kinase subunit of the insulin growth factor pathway and has been implicated in starvation resistance in a cancer study [62]. Aside from insulin signaling, the top two genes shared between methods have been implicated in starvation resistance. Epidermal growth factor receptor (*Egfr* ) is important for normal cell growth, while overexpression of the receptor is a common route for cancer development [63]. Sevelda *et al*. showed that *Egfr* increased starvation resistance through interactions with *Akt1* [64]. *Erk7* is an extracellular kinase involved in the secretory system. *Erk7* downregulates secretion by triggering the destruction of endoplasmic reticulum exit sites under starvation conditions [65].

Most of the genes found by GO-BayesC only are from the nucleoporin family (*mbo, Nup53, Nup54, Nup93-1, Nup93-2, Nup98-96, Nup153, Nup205* ). Nucleoporin degradation can occur under starvation conditions to prevent the export of macromolecules [53]. The remaining genes are tangential to the *InR* signaling pathway. The adipokinetic hormone (*Akh*) makes a complex with its receptor *AkhR* previously described as part of the *InR* signaling pathway. Downstream in the pathway, a signal propagating kinase *Pi3K92E* was also found by GO-BayesC. Eleven distinct genes found by GO-TBLUP only are involved in various complexes and pathways. Three of these genes are related to the Hippo, or Salvador-Warts-Hippo, signaling pathway [66] — the core signaling kinase hippo (*hpo*), a kinase further downstream (*ack* ), and *Sik3. Sik3* is involved in balancing NADPH/NADP+, a set of key components for oxidative stress mitigation and cellular signaling [67]. *Sik3* is also part of the *InR* signaling pathway that negatively regulates Hippo signaling. Another top gene (*pdk1* ) is part of the *InR* signaling pathway and is responsible for inhibiting apoptosis during embryonic development. In flies, activin signaling responds to carbohydrate levels in the gut by increasing carbohydrase expression [68]. Additionally, activin signaling increases starvation resistance for neuronal cells [69]. The products of two genes, *put* and *babo*, form a complex with an activin-like ligand to initiate the activin signaling pathway [70]. Hemipterous (*hep*) is a kinase involved in imaginal disc formation and cell proliferation. A study found that *hep* loss-of-function mutant flies have reduced energy stores and lower starvation resistance. [71] *hep* has also been implicated in obesity and insulin resistance by activating the JNK signaling pathway [72].

In males, GO-BayesC and GO-TBLUP found eight genes in common out of the top 1% of GO terms. The two major categories that emerge from these genes are carbohydrate metabolism and cell polarization. For carbohydrate metabolism, the insulin-like receptor (*InR*) and an insulin-like peptide (*Ilp2* ) are key components of *InR* signaling. *InR* is the only gene that was found by both GO methods in both sexes. The remaining genes are all related to cell polarization. The Crumbs complex has three genes (*crb, sdt, Patj* ), while the PAR complex has two genes (*aPKC* and *par-6* ) from both methods. Both complexes are highly conserved regulators of apico-basal cell formation [73]. The PAR complex negatively regulates the *InR* signaling pathway [74]. Outside of these complexes, *PDZ-GEF* is a guanine exchange factor involved in epithelial cell polarization through a separate mechanism [75].

Six genes were found by GO-BayesC only in the top 1% of GO terms from males. Three genes are related to the PAR complex. Bazooka(*baz* ) is a core component of the PAR complex while two genes, *scrib* and *dlg1*, form a conserved complex with *lgl* that regulates cell migration with the PAR complex [76]. The product of *Moe* competes with PAR complex component *aPKC* to regulate the Crumbs complex [77]. *AkhR* was found uniquely by GO-BayesC. As previously described, this gene is involved in carbohydrate and lipid metabolism signaling processes [61]. Another gene, shotgun(*shg* ), is downstream in the *Egfr* signaling pathway [78]. As previously described, *Egfr* promotes starvation resistance through *Akt1* interactions [64]. Only two genes were found uniquely by GO-TBLUP in males. One is a desaturase (*Desat1* ) that synthesizes fatty acid molecules. *Desat1* has been shown to induce cell autophagy under starvation conditions [79]. The other, *Ras85D*, is an oncogenic cell growth promoter. Downregulating *Ras85D* has been shown to improve starvation resistance by limiting growth signals [80].

## Discussion

In this study, we evaluated ten statistical methods on their ability to predict starvation resistance, a well-documented quantitative complex trait [21], using transcriptomic data [22]. As expected, we found differences in the prediction accuracy provided by methods tested, both within sex and between sexes. While most methods were somewhat predictive, neural networks provided minimal prediction accuracy for both sexes. This is in agreement with previous work showing the importance of feature selection prior to model fitting in the *n << p* regime for neural networks to perform well [11]. However, here we wanted to focus on out-of-the-box performance of the different methods. The most predictive methods in females, VARBVS and MR.ASH, are both Bayesian regression methods that perform effect shrinkage and variable selection. These methods allow for the underlying genetic architecture to be sparse, suggesting that not all genes affect starvation resistance. In contrast, the most predictive methods in males, TBLUP and ridge regression, only perform effect shrinkage, which suggests that the genetic architecture of starvation resistance is denser in males. The difference in best performing methods between sexes is not surprising because starvation resistance is a known sexually dimorphic trait with different genetic architectures between sexes [33, 81]. This difference highlights the importance of choosing methods with assumptions that match the complex trait under investigation. On the other hand, prediction analysis can also provide some hypothesis about the genetic architecture of the trait analyzed, to be followed up with more specific experiments.

Previous studies have shown that including functional annotation information into prediction models can help improve predictions [17, 45–47]. Thus, we selected two methods that allow the incorporation of additional information, BayesC and TBLUP, and annotated them with Gene Ontology (GO) information. We showed that a small number of biologically relevant GO terms achieved substantially higher prediction accuracies for GO-BayesC and GO-TBLUP than the standard BayesC and TBLUP. While the correlation between prediction accuracies for each GO between GO-BayesC and GO-TBLUP was high for both sexes (*r* = 0.84), differences in method characteristics resulted in different top GO terms between methods, especially in males. The most predictive GO terms for both sexes shared genes such as *InR* and *AkhR* that are involved in carbohydrate and/or lipid metabolism. Both genes have been associated to starvation resistance in previous studies [60, 61]. However, many of the genes shared by the most predictive GO terms in the two sexes were different (*e*.*g*., *Egfr* in females and *sdt* in males). These findings also suggest that both methods may be able to highlight sex-shared and sex-specific genes of interest for traits with unknown genetic architectures. Overall, our findings suggest that additional information from GO terms may help disentangle signal from noise to improve prediction accuracy and our understanding of the complex trait of interest.

In conclusion, we found that differences in prediction performance between methods exists and depend on the assumptions made by the model relative to the trait of interest. We also confirmed that external information such as GO term annotation can improve prediction accuracy for biologically relevant data. However, there are a number of limitations and considerations to address. First, the data is limited by the small number of available DGRP lines. Each transcriptome containing over 11,000 genes results in a *n << p* problem that makes estimation of the relevant parameters problematic. We hypothesize that all methods would improve by increasing the sample size of the DGRP. Second, the linear regressions performed by most methods used here do not account for non-linear, or epistatic, interaction effects between genes and may perform poorly for complex traits with epistatic effects. Third, the gene expression data were obtained from whole flies under standard conditions. Higher prediction accuracy may be achieved if gene expression were measured under the same starvation conditions used to score starvation resistance, and for relevant tissues. Despite these limitations, we showed that gene expression data coupled with appropriate model selection can be effective for complex trait prediction.

## Supporting information

**S1 Fig**. Violin plot comparison of Sparse Group Lasso results for top GO terms from GO-BayesC/GO-TBLUP along with randomly selected GO terms in females.

**S2 Fig**. Distribution of number of overlapping genes for top 1% of GO terms for GO-BayesC and GO-TBLUP in females and males. The selection cutoff is marked by the vertical bar.

**S1 Table**. Mean prediction accuracy and standard error for all methods in females and males.

**S2 Table**. Mean prediction accuracy and standard error for each GO term in GO-BayesC and GO-TBLUP for females and males.

**S3 Table**. Genes in the top 1% of GO terms for GO-BayesC and GO-TBLUP in females and males ordered by gene count.

## Data and code availability

All DGRP lines are available from the Bloomington Drosophila Stock Center (Bloomington, IN). All raw and processed RNA-Seq data are available at the NCBI Gene Expression Omnibus (GEO; https://www.ncbi.nlm.nih.gov/geo/) under accession number GSE117850. Phenotypic data are available at http://dgrp2.gnets.ncsu.edu/. The code used for the analyses is available at https://github.com/nklimko/dgrp-starve.

## Acknowledgments

We thank Trudy Mackay for helpful comments on an earlier version of this manuscript, and Liangjiang Wang for suggestions about the neural network analyses. Research reported in this publication was in part supported by the National Institute of General Medical Sciences of the National Institutes of Health under Award Number R35GM146868 to FM. The content is solely the responsibility of the authors and does not necessarily represent the official views of the National Institutes of Health.

