## Supplementary figures and images for "Comparing statistical learning methods for complex trait prediction from gene expression"

### S1 Fig

A

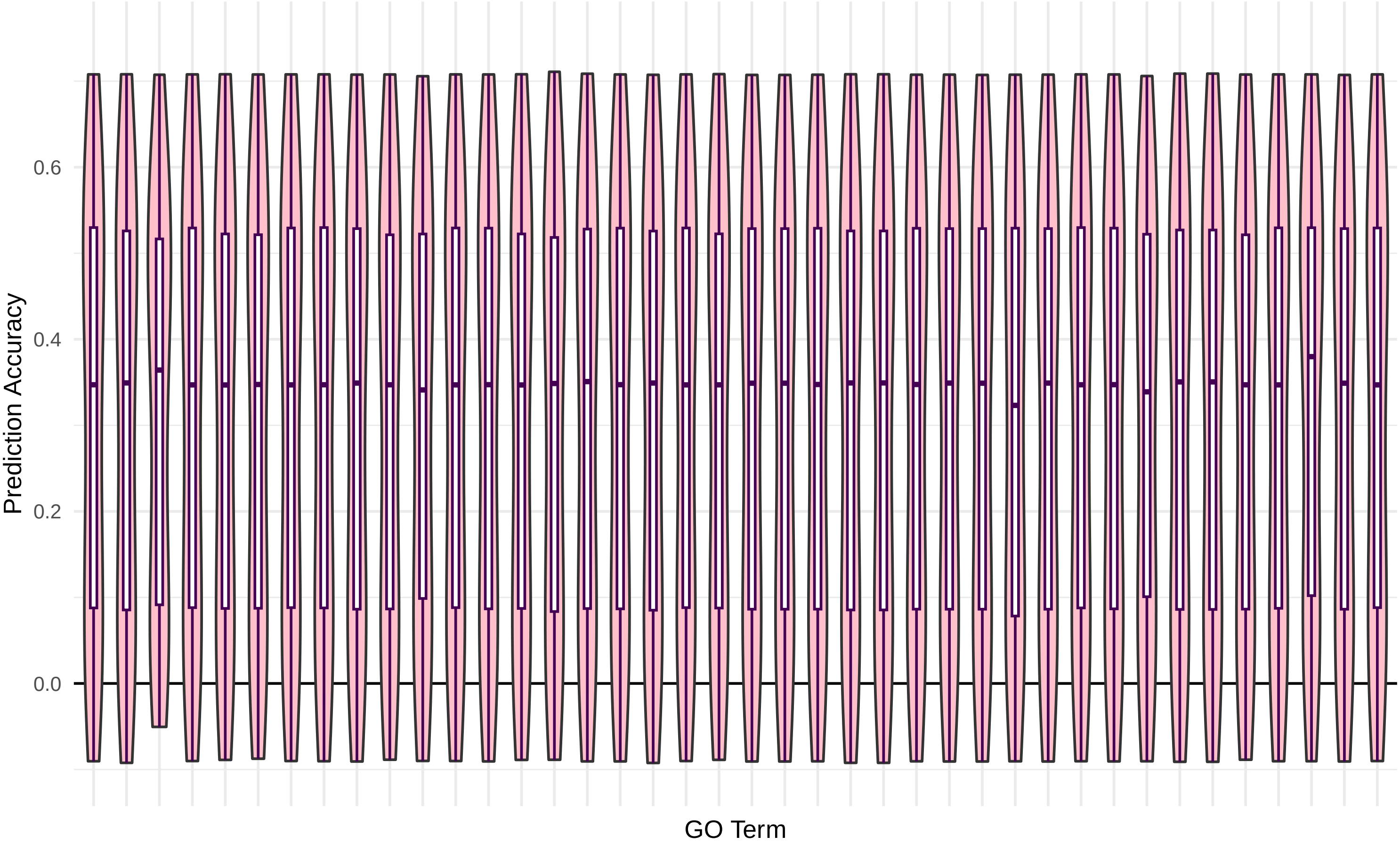

### S2 Fig

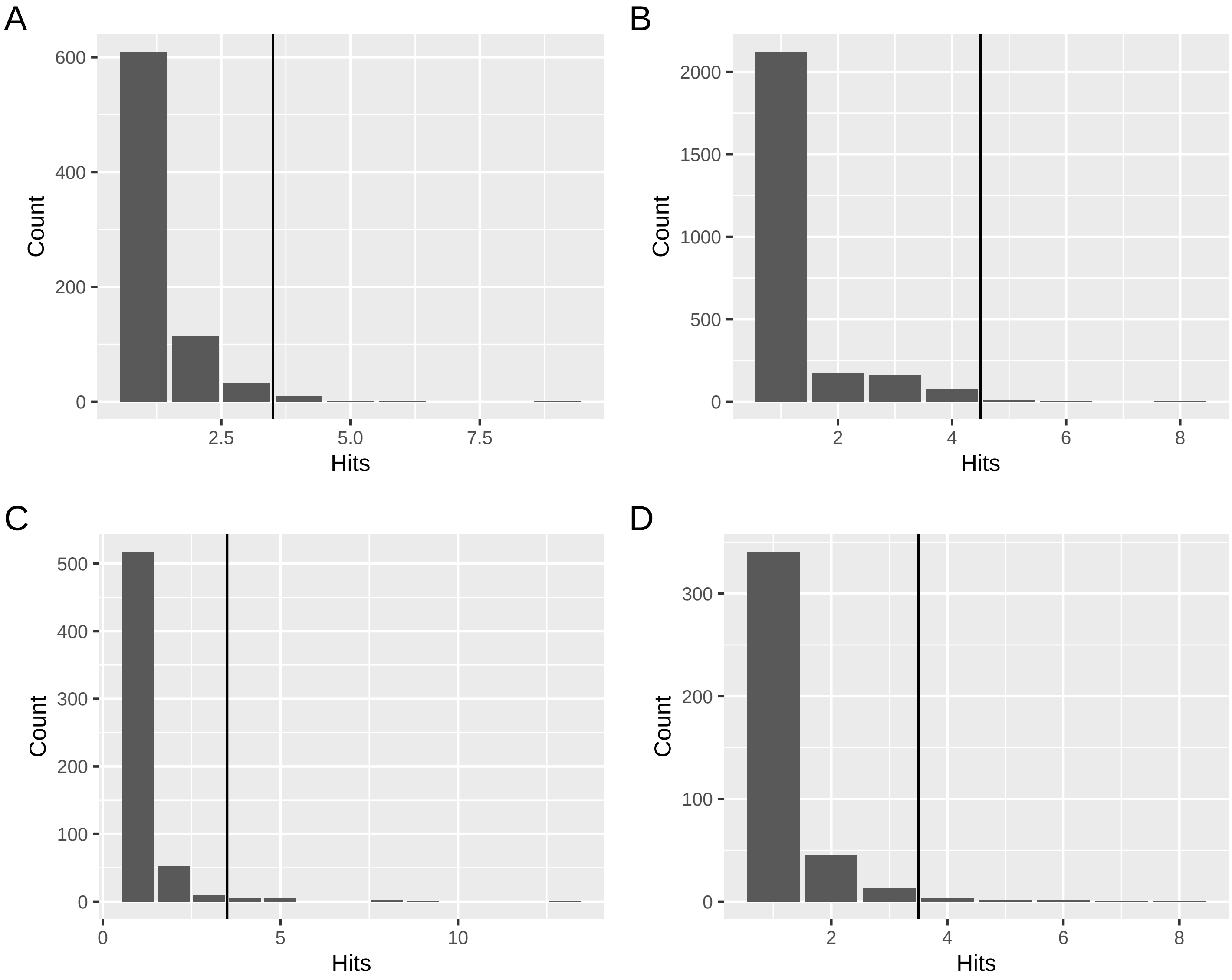
